# Development of a low-dose PBMC humanized mouse model using CD47;Rag2;IL2rγ triple KO mice: Enhanced leukocyte reconstitution and extended experimental window

**DOI:** 10.64898/2026.03.25.714298

**Authors:** Kang-Hyun Kim, Hye-Young Song, Sang-wook Lee, In-Jeoung Baek, Je-Won Ryu, Seon-Ju Jo, Seung-Hee Ryu, Sun-Min Seo, Seung-Ho Heo

**Affiliations:** Convergence Medicine Research Center, Asan Medical Center, Seoul, Republic of Korea; College of Veterinary Medicine, Chonnam National University, Gwangju, Republic of Korea; Asan Institute for Lifesciences, Asan Medical Center, Seoul, Republic of Korea; Department of Radiation Oncology, Asan Medical Center, Seoul, Republic of Korea; Laboratory Animal Research Center, Konkuk University, Seoul, Republic of Korea

## Abstract

Humanized mice (hu-mice), which recapitulate the human immune system, have become increasingly important for preclinical immunotherapy studies. Among these models, the human peripheral blood mononuclear cells (PBMC)-engrafted hu-mice model is the simplest and fastest. However, its utility is hindered by the development of lethal graft-versus-host disease (GvHD) and the insufficient reconstitution of human leukocytes. To address these limitations, we developed PBMC hu-mice models using a novel strain, NOD-CD47^null^Rag2^null^IL-2rγ^null^ (RTKO) focusing on the immunological defects of the NOD strain and the immunotolerance provided by CD47 deficiency. Six-week-old female NOD-Rag2^null^IL-2rγ^null^ (RID) and RTKO mice were intravenously injected with three different PBMC doses (3×10^6^, 5×10^6^, and 1×10^7^ cells). At standard doses (5×10^6^ and 1×10^7^ cells), RTKO mice exhibited enhanced engraftment of human leukocytes, though GvHD was more severe compared to the RID strain, resulting in a limited experimental window. However, in a subsequent trial using a lower dose of PBMCs (3 × 10^6^ cells), RTKO mice demonstrated notable advantages, including stable reconstitution of human leukocytes, milder GvHD symptoms without life-threatening lesions, and a markedly prolonged experimental window. Considering the difficulties in generating hematopoietic stem cell (HSC)-engrafted hu-mice, the extended experimental window provided by this model, which is comparable to HSC hu-mice, is a significant improvement. Moreover, the radiation tolerance conferred by the Rag gene mutation in this model offers another advantage for radiotherapy research. Consequently, the low-dose PBMC RTKO model serves as a versatile and valuable platform for a broad spectrum of immunotherapy studies, especially in the field of immuno-oncology.

## Introduction

The development of animal models that closely mimic human biological systems continues to evolve, playing a significant role in both basic research and preclinical trials to develop treatments. Among these, humanized mice (hu-mice), which recapitulate a functional human immune system, are highly desirable in the field of immunotherapy [1–3]. Since the late 1980s, hu-mice have been generated by transplanting human cells or immune tissues into immunodeficient mice. The success of producing hu-mice depends on selecting an appropriate immunodeficient mouse strain and implementing an efficient transplantation protocol [4, 5].

The most commonly used immunodeficient mouse strains for generating hu-mice are NOD-SCID;IL2rγ^null^ (NSG or NOG) and NOD-Rag^null^IL2rγ^null^ (NRG or RID). Mutations in the *Prkdc* (also known as XRCC7) gene or the knockout of recombination-activating genes (RAG) in mice lead to deficiencies in T and B cells [2, 6]. Both NSG and NRG mice are widely used in the generation of hu-mice [6], but those with the Rag gene mutation offer advantages such as tolerance to irradiation and resistance to T/B cell leakiness [7]. Knocking out the interleukin-2 receptor gamma chain (IL2rγ) prevents the development of natural killer (NK) and NKT cells, promoting phagocytic tolerance to xenografts and improving the engraftment of human immune cells [4, 8]. In addition, IL2rγ deficiency disrupts signaling of various cytokines, including IL-2, IL-4, IL-7, IL-9, IL-15, and IL-21 [9, 10]. Moreover, the non-obese diabetic (NOD) strain has the advantages of functional defects in macrophages and dendritic cells, reduced complement activity, and polymorphisms in signal regulatory protein α (SIRPα), compared to other mouse strains such as Balb/c and C57BL/6 [11–13].

Humanized mouse models are categorized into three representative types based on the transplanted immune cells or tissues: (a) human peripheral blood mononuclear cells (PBMCs); (b) CD34+ hematopoietic stem cells (HSCs); and (c) bone marrow, liver, and thymus (BLT) (3, 4). Each model has its own advantages and limitations. Among these, the PBMC-engrafted hu-mice model is known as the simplest, fastest, and most cost-effective approach due to the relatively easy availability of PBMCs, no requirement for myelosuppression, and the reconstitution of human immune cells within two to three weeks. However, this model has drawbacks compared to other models, including a high incidence of severe, lethal graft-versus-host disease (GvHD), which results in a narrow experimental window of only three to six weeks. Furthermore, the immune reconstitution is largely limited to human T cells, with minimal representation of other hematopoietic lineages [2, 6, 14]. To overcome these limitations, ongoing efforts have focused on improving the PBMC hu-mice model by modifying host immune genes [15, 16], inserting human genes [17–19], and administering human cytokines to the mouse hosts [6, 20].

CD47, a ubiquitously expressed protein in the immunoglobulin superfamily, serves as a ligand for signal regulatory protein α (SIRPα). SIRPα, another immunoglobulin superfamily protein, is abundant in myeloid cells, particularly macrophages and leukocytes. The CD47-SIRPα signaling system inhibits host cell phagocytosis, with CD47 functioning as a “don’t eat me” signal [21–23]. Focusing on this function, humanized mouse models have been developed using CD47-deficient strains [16, 24, 25]. However, models using the NOD background, which express SIRPα with high affinity for human CD47 and exhibit functional immunodeficiencies in macrophages and dendritic cells, are still relatively rare [26, 27]. In this study, we generated PBMC hu-mice using CD47;Rag2;IL2rγ triple KO NOD (RTKO) mice, aiming to enhance the engraftment of human immune cells and alleviate GvHD symptoms. We applied three PBMC doses to optimize the conditions for hu-mice generation and presented an improved model that overcomes the existing limitations, based on clinical symptoms, immunological assessments, and histopathological analyses.

## Materials and Methods

### Animals

This study strictly complied with the guidelines set by the Institutional Animal Care and Use Committee of the Asan Institute of Life Sciences (Seoul, Korea, IACUC No 2022-12-111). All animals were housed under specific pathogen-free conditions and bred in individually ventilated cages (Blue line; Tecniplast S.p.A., VA, Italy). Irradiated rodent chow (Teklad rodent diets 2018; Inotiv, Inc., IN, USA) and reverse-osmosis water were provided ad libitum. Rag2;IL2rγ double KO NOD (RID) mice were purchased from GEM Biosciences (Cheongju, Korea). CD47;Rag2;IL2rγ triple KO NOD (RTKO) mice were generated by crossing RID mice with CD47 KO NOD mice provided by Dr. In-Jeoung Baek from the Convergence Medicine Research Center in Seoul, Korea. The genotypes for CD47, Rag2, and IL2rγ were examined as previously described [27].

### Experimental design

Six-week-old female RID and RTKO mice were used to generate PBMC-humanized mice. PBMCs purchased from Lonza (4W-270C; Basel, Switzerland) were thawed with RPMI 1640 medium (Thermo Fisher Scientific, Waltham, MA, USA) supplemented with 10% FBS and 1% penicillin/streptomycin. After then, PBMCs in three different doses—1 × 10^7^ (high dose; HD), 5 × 10^6^ (middle dose; MD), and 3 × 10^6^ (low dose; LD)—were suspended in 300 μL of phosphate-buffered saline and immediately injected intravenously into RID and RTKO mice. We used PBMCs from the same donor between RID and RTKO mice (the MD and HD groups, batch no. 3038065; the LD groups, batch no. 3038103). To prevent bacterial infection, enrofloxacin (0.27 mg/mL; Bayer, Leverkusen, Germany) was administered through drinking water [27]. Clinical signs were monitored twice a week, and body weight was measured once a week. Starting from two weeks post-transplantation (wpt), 100 μL of blood was collected from the retro-orbital plexus every week following inhalation anesthesia using 2% isoflurane (Terrel™, Piramal Critical Care, Inc., Bethlehem, PA, USA). The HD and MD groups were euthanized at 6 wpt, while the LD group was euthanized at 14 wpt. At the time of the euthanasia, mice were administered 50 mg/kg of alfaxan (Jurox, Rutherford, Australia) and 10 mg/kg of xylazine (Rompun™; Elanco, Ansan, Korea) via intraperitoneal injection for general anesthesia [28]. Once fully anesthetized, the mice were sacrificed by exsanguination via the inferior vena cava for further analysis.

### Flow cytometry analysis

Peripheral blood samples were collected from the retro-orbital plexus using heparin-coated capillary tubes (Paul Marienfeld GmbH & Co.KG, Lauda-Königshofen, Germany). Red blood cells were lysed by incubating the samples with RBC lysis buffer (Biolegend, San Diego, CA, USA) for 20 minutes. And then, the cells were washed with Dulbecco’s PBS (DPBS), Fc binding was blocked for 5 minutes at 4°C using human BD Fc Block (Becton Dickinson Biosciences, Franklin Lakes, NJ, USA), and stained using conjugated antibodies. The antibodies used in this study were: anti-hCD45-pacific blue (HI30), anti-hCD3-APC-Cy7 (UCHT1), anti-hCD19-PerCP-cy5.5 (HIB19), anti-hCD4-FITC (RPA-T4), anti-hCD8-PE (RPA-P8), and anti-hCD56-Amcyan (NCAM16.2) from BD Biosciences, anti-hCD66b-PE-Cy7 (G10F5), and anti-hCD14-APC (HCD14) from Biolegend. Data were acquired using a FACS Canto Ⅱ Flow cytometer (BD Biosciences) and analyzed using FACSDiva 8.0.2 software (BD Biosciences).

### Hematoxylin and eosin (H&E) staining and histopathological analysis

Fixed lung, liver, kidney, and skin tissues were processed using standard methods, embedded in paraffin, and then cut into 4-μm sections. The sections were deparaffinized, rehydrated, and stained with hematoxylin and eosin (H&E). After staining, the sections were dehydrated, cleared, mounted, and examined under light microscopy. A semi-quantitative scoring system (ranging from 0 to 5 grades) was used to assess the severity of the lesions as follows [19, 20]: (0) Normal; (1) Minimal: minimal inflammatory cell aggregation; (2) Mild: inflammatory cell aggregation (≤ 10%), minimal apoptosis or necrosis, and epithelial thickening (≤ 30 µm); (3) Moderate: inflammatory cell aggregation (≤ 25%), apoptosis or necrosis, and epithelial thickening (≤ 60 µm); (4) Severe: inflammatory cell aggregation (≤ 50%), necrotic foci, and epithelial thickening (≤ 100 µm); (5) Generalized: inflammatory cell aggregation (≥ 50%), necrotic foci, and epithelial thickening (≥ 100 µm). Two veterinary pathologists independently reviewed all the lesions.

### Immunohistochemical staining

For immunohistochemistry, selected serial sections (4 μm) were deparaffinized, rehydrated, and placed in 0.01 M citrate buffer (pH 6.0). The sections were then heated in a microwave for 20 minutes (two 10-minute intervals, with a 10-minute break in between). Then, the slides were incubated for 15 minutes in 1.0% H_2_O_2_. The slides were preincubated with blocking serum (Vectastain ABC kit; Vector Laboratories, Burlingame, CA, USA), followed by incubation with rat anti-human CD45 (MA5-17687, 1:100, Invitrogen), mouse anti-human CD19 (14-0199-82; 1:150, Invitrogen), and mouse anti-human CD3 antibodies (14-0038-82; 1:100, Invitrogen). The sections were then incubated with biotinylated secondary antibodies and then with avidin-coupled peroxidase using the Vectastain ABC kit (Vector Laboratories). The CD3 and CD19 antibodies were stained using the Mouse on Mouse detection kit (Vector Laboratories). After development with 3,3′-diaminobenzidine (DAB) substrate (DAB Substrate kit, Vector Laboratories), the slides were counterstained with hematoxylin. Positive cells were manually counted within a 0.03 mm^2^ area under a light microscope. Statistical analysis was performed based on counts obtained from three representative fields per slide.

### Human cytokine measurement

At the time of sacrifice, whole blood was collected from the inferior vena cava of each mouse under general anesthesia. The blood was centrifuged for 10 min at 10,000 ×g, and the resulting serum samples were stored at −80°C for further analysis. Each serum sample was analyzed for human cytokines using the Human CorPlex™ Cytokine Panel 1 10-Plex Array (116-7BF-1-AB; Quanterix, Billerica, MA, USA) according to the manufacturer’s instructions. The cytokines measured included human IL-1β, IL-4, IL-5, IL-6, IL-8, IL-10, IL-12p70, IL-22, IFN-γ, and TNF-α. The results were acquired and analyzed using the SP-X Imaging System (Quanterix).

### Statistical analysis

Survival curves were plotted using Kaplan–Meier estimates. Statistically significant differences between the groups were evaluated using the unpaired Student’s t-test or Fisher’s exact test. All statistical analyses were performed using GraphPad V8.0.2 software (GraphPad Software Inc., Billerica, MA, USA). All data are expressed as the mean ± standard deviation (SD), with statistically significant results indicated by asterisks (*) on the graphs. * P < 0.05, ** P < 0.01

## Results

### Generation of middle- and high-dose PBMC hu-mice and their clinical signs

Six-week-old female RID and RTKO mice were used for generating PBMC Hu-mice. Clinical signs and overall animal health were monitored twice a week, and body weight was measured once a week. The observed clinical symptoms of GvHD included hyperkeratosis, anemia, jaundice, and weight loss. The animal experiments were concluded at 6 wpt based on the health status and survival rate of the mice. At the study endpoint, the final survival rates were as follows: RID-MD 83.3% (n = 6), RID-HD 66.7% (n = 6), RTKO-MD 85.7% (n = 7), and RTKO-HD 57.1% (n = 7). Although the RTKO-HD group exhibited the lowest survival rate, no statistically significant differences were observed between the groups (Fig 1A). Detailed information on animal mortality is provided in Table 1. Body weight began to decline at 4 wpt in the MD groups and at 3 wpt in the HD groups. At necropsy, all groups except the RID-MD group showed decreased weight compared to their initial measurements. The body weight ratio of the RTKO-HD group was significantly lower than that of the RID-MD group (Fig 1B).

**Fig 1.**
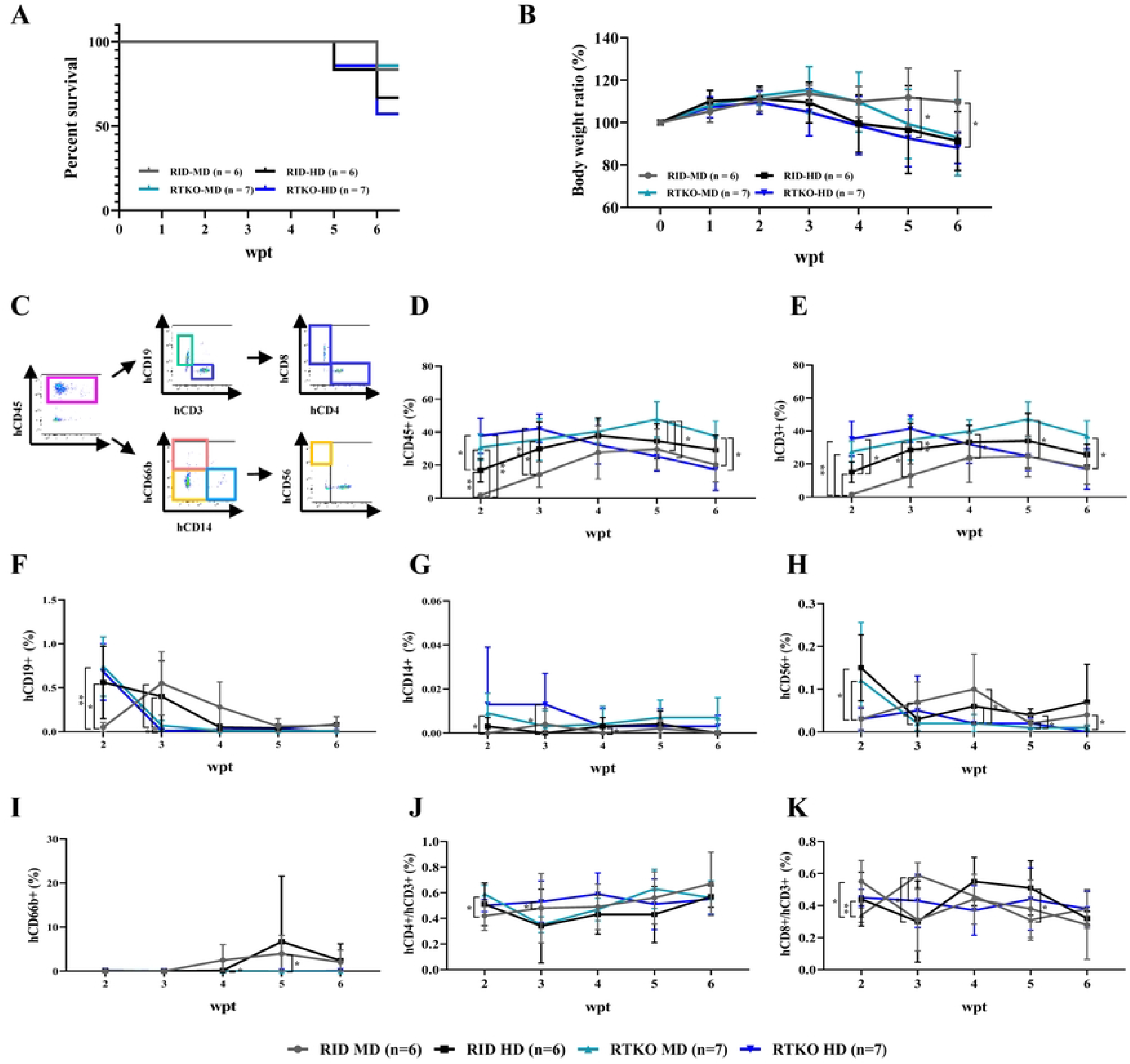
Clinical observation and immune monitoring of PBMC-engrafted humanized mice models using standard doses. (A) Survival rates and (B) body weight ratios (calculated as a percentage of the original body weight measured at the time of PBMCs infusion) were monitored weekly after PBMCs transplantation. The engraftment of human cells was assessed via FACS analysis weekly from two to six weeks post-transplantation. (C) Gating strategy for flow cytometry analysis. Serial analysis of (D) hCD45+ leukocytes, (E) hCD3+ T cells, (F) hCD19+ B cells, (G) hCD14+ monocytes, (H) hCD56+ NK cells, and (I) hCD66b+ granulocytes were examined. Within hCD3+ cells (considered 100%), the percentages of (J) hCD4+ cells and (K) hCD8+ cells were calculated. Representative dot plots for hCD45, hCD3 and hCD19, hCD4 and hCD8, and hCD14 and hCD66b are provided in S1 Fig. The number of mice in each experimental group was as follows: RID-MD (n = 5-6), RID-HD (n = 4-6), RTKO-MD (n = 6-7), RTKO-HD (n = 4-7). *P < 0.05 and **P < 0.01. PBMC, peripheral blood mononuclear cell; RID, Rag2; IL2rg double KO NOD mice; RTKO, CD47; Rag2; IL2rg triple KO NOD mice; MD, middle dose; HD, high dose; FACS, fluorescence-activated cell sorting; wpt, weeks post transplantation

**Table 1.** Detailed information on animal mortality.

| Groups | Number and timing of deaths | Clinical signs or necropsy findings |
| --- | --- | --- |
| RID-LD<br>(n = 7) | Two euthanized mouse at 9 wpt | Weight loss of 18.1%, jaundice, moderate hepatic inflammation<br>No weight loss, mild anemia, hypothermia, moderate pulmonary inflammation |
| RID-MD<br>(n = 6) | One dead mouse at 6 wpt | Sudden death, no weight loss, mild pulmonary inflammation, and splenomegaly |
| RID-HD<br>(n = 6) | One euthanized mouse at 5 wpt | Weight loss of 22.1%, anemia, mild splenomegaly |
|  | One euthanized mouse at 6 wpt | Weight loss of 31.2%, jaundice, moderate hyperkeratosis, and hepatic inflammation |
| RTKO-LD<br>(n = 8) | One dead mouse at 6 wpt | Sudden death, weight loss of 18.1%, anemia, mild pulmonary inflammation, and splenomegaly |
|  | One dead mouse at 12 wpt | Sudden death, weight loss of 6.0%, moderate hyperkeratosis, mild hepatic inflammation |
| RTKO-MD<br>(n = 7) | One euthanized mouse at 6 wpt | Weight loss of 25.5%, mild hyperkeratosis, moderate pulmonary inflammation, and splenomegaly |
| RTKO-HD<br>(n = 7) | One euthanized mouse at 5 wpt | Weight loss of 21.7%, anemia, moderate hepatic inflammation |
|  | One euthanized mouse at 6 wpt | Weight loss of 14.4%, reduced activity, mild hepatic and pulmonary inflammation |
|  | One dead mouse at 6 wpt | Sudden death, weight loss of 21.0%, mild hyperkeratosis, moderate hepatic inflammation |

RID, Rag2; IL-2 $\gamma$ double KO NOD mice; RTKO, CD47; Rag2; IL-2 $\gamma$ triple KO NOD mice; LD, low dose; MD, middle dose; HD, high dose; wpt, weeks post-transplantation.

### Enhanced human immune cell engraftment in RTKO mice

To assess human immune cell transplantation, flow cytometry for human leukocyte antigens was performed weekly from 2 wpt. The gating strategy used for flow cytometry analysis is depicted in Figure 1C. At 2 wpt, more than 10% engraftment of human leukocytes was observed in all groups except the RID-MD group. The hCD45 ratios in the RTKO groups exceeded 30%, which was more than double the levels observed in the RID groups. The engraftment of human immune cells initially increased and then gradually decreased. The RTKO-HD group, which exhibited the highest hCD45 ratio at the first measurement, showed a decrease from 4 wpt. The hCD45 ratio in the RID-HD group began decreasing at 5 wpt, while the remaining groups showed declines at 6 wpt. The RTKO-MD group exhibited the peak hCD45 ratio at 5 wpt (47.4%), which was significantly higher compared to those of all other groups. At the final measurement, the RTKO-MD group had the highest hCD45 ratio, while the RTKO-HD group had the lowest ratio (Fig 1D). The ratios of hCD3 followed a similar kinetic pattern to those of hCD45, indicating that the majority of the engrafted human immune cells were T cells (Fig 1E). Overall, hCD4 ratios increased and hCD8 ratios decreased over time, though no consistent differences were observed between the groups (Figs 1J and 1K). The ratios of hCD14, hCD19, and hCD56 remained low at less than 1%, with hCD56 levels being higher in the RID groups and those of hCD14 elevated in the RTKO group (Figs 1F–1H). Some RID mice showed significant levels of hCD66b at 5 wpt (RID-MD 3.97%, RID-HD 6.69%) (Fig 1I). Dot plots for hCD45, hCD3 and hCD19, hCD4 and hCD8, hCD14 and hCD66b, and hCD56 are provided in S1 Fig.

### Alleviation of GvHD symptoms and the extended experimental window in low-dose PBMC hu-mice

The MD and HD PBMC hu-mice exhibited a limited experimental period of 6 weeks, primarily due to the onset of lethal GvHD symptoms. To address this limitation, we developed a low-dose PBMC hu-mice model by administering 3 × 10^6^ PBMCs. The body weight of the RID-LD group continued to increase until 13 wpt, and that of the RTKO-LD group decreased after 11 wpt; however, no significant difference was observed between the two groups. Comparing the final body weight ratios across all groups, the RID-LD and RTKO-LD groups exhibited significantly higher ratios, with a decreasing trend observed as the PBMC dosage increased (Fig 2B). In the RID-LD group, two mice were euthanized at 9 wpt owing to jaundice and severe hypothermia, while in the RTKO-LD group, one mouse died at 6 wpt and another at 12 wpt. Detailed information on animal mortality is provided in Table 1. The final survival rates at 14 wpt were 71.4% for the RID-LD group (N = 7) and 75.0% for the RTKO-LD group (N = 8), with no significant differences across all groups (Fig 2A). At the end of the experiment, no specific clinical or necropsy findings were observed in the RID-LD group, except for hyperkeratosis in some mice. In the RTKO groups, only two mice with reduced body weight showed mild atrophy and inflammation in the liver, lung, and kidney. In the low-dose PBMC hu-mice, the reconstitution of human immune cells was delayed, with meaningful hCD45 levels observed from 3 wpt. The hCD45 ratio in the RTKO-LD group steadily increased, reaching the peak value (36.8%) at 7 wpt and then gradually decreasing to 23.3% at 14 wpt. Since the human leukocytes ratio was maintained above 10% from 5 to 14 wpt, the stable experimental period can be considered approximately 10 weeks (Fig 2C). In the RID-LD group, the hCD45 ratio was above 10% only for three weeks (6 to 8 wpt), reaching a maximum of 21.9% at 8 wpt before rapidly dropping to below 10% after the euthanasia of two mice (Fig 2C). Individual hCD45 ratios for the LD groups are provided in S1 Table. Comparison of the maximum hCD45 levels across all groups revealed that the LD groups exhibited the lowest values within each mouse strain, although there were no meaning differences. However, the values of the RID-LD and MD groups were significantly lower than those of the RTKO-MD group and the final values were significantly lower in the LD groups. Most of the transplanted human leukocytes were hCD3+ T cells, and their proportion was almost identical to that of hCD45 (Figs 2C and 2D). There were no significant differences in the hCD4 and hCD8 ratios in the hCD3+ population (Figs 2G and 2H). Significant engraftment of hCD19+ B cells and hCD66b+ leukocytes was observed in the RTKO-LD group compared to the RID-LD group; however, the ratio was extremely low, at less than 0.2%, and significantly lower than that observed in the MD and HD groups (Figs 2E and 2F). hCD14+ monocytes and hCD56+ NK cells were barely detectable (Data not shown). Dot plots for hCD45, hCD3 and hCD19, hCD4 and hCD8, and hCD14 and hCD66b are provided in S2 Fig.

**Fig 2.**
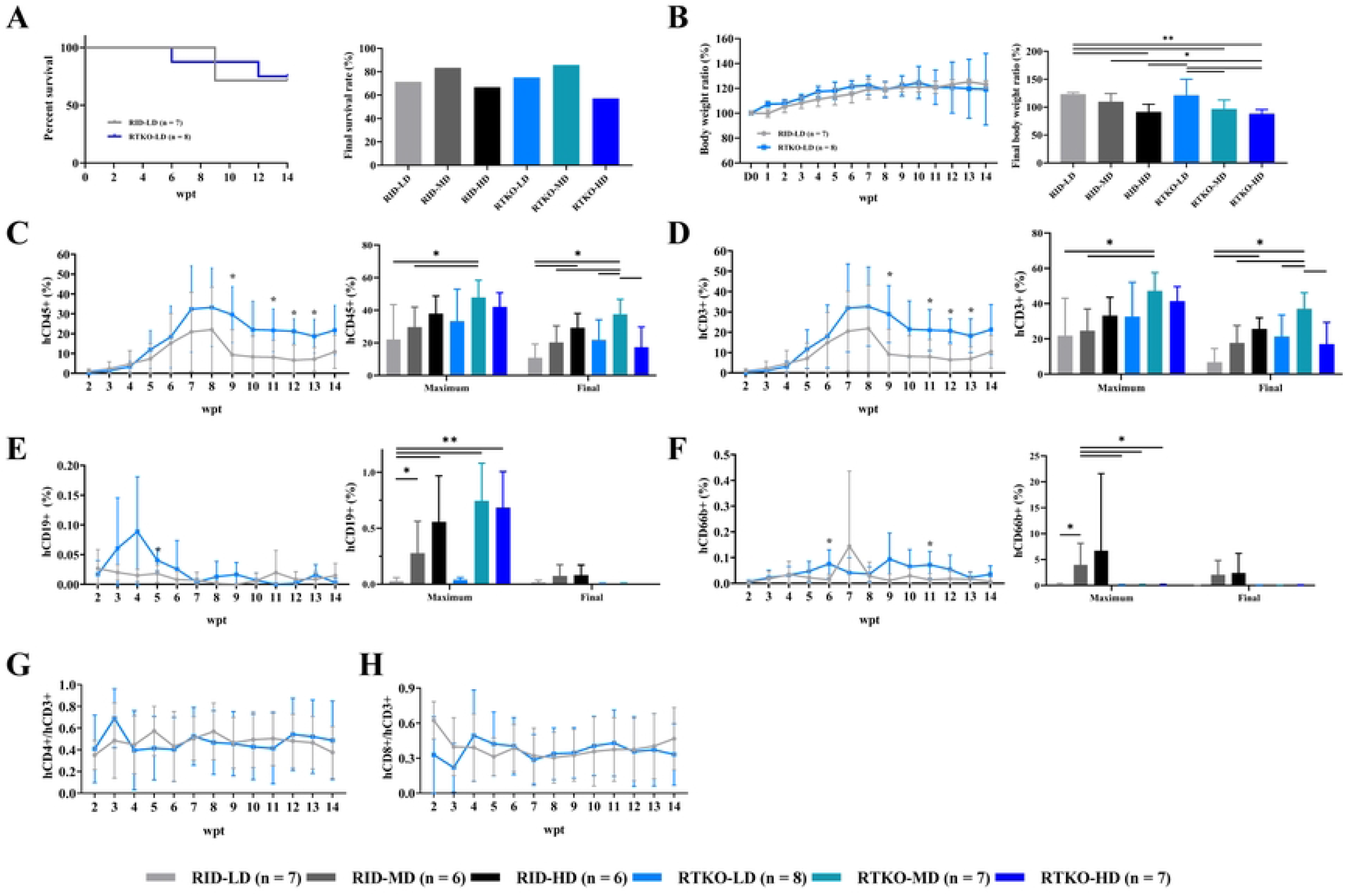
Clinical observation and immune monitoring of a lower dose of PBMC-engrafted humanized mice model and comparative analysis across all groups. (A) Survival rates and (B) body weight ratios (calculated as a percentage of original body weight measured at the time of PBMCs transplantation) for the LD groups, alongside the final outcomes for all experimental groups. The engraftment of human cells was assessed via FACS analysis weekly from 2 to 14 weeks after PBMCs injection. Serial measurements of (C) hCD45^+^ leukocytes, (D) hCD3^+^ T cells, (E) hCD19^+^ B cells, and (F) hCD66b^+^ granulocytes in the LD groups, along with the peak and final results for each group. Within hCD3+ cells (considered 100%), the percentages of (G) hCD4+ cells and (H) hCD8+ cells were calculated. Representative dot plots for human leukocyte antigens of the LD groups are presented in S2 Fig. The number of mice in each experimental group was as follows: RID-LD (n = 5-7), RID-MD (n = 5-6), RID-HD (n = 4-6), RTKO-LD (n = 6-8), RTKO-MD (n = 6-7), RTKO-HD (n = 4-7). *P < 0.05 and **P < 0.01. PBMC, peripheral blood mononuclear cell; GvHD, graft-versus-host disease; RID, Rag2; IL2rg double KO NOD mice; RTKO, CD47; Rag2; IL2rg triple KO NOD mice; LD, low dose; MD, middle dose; HD, high dose; FACs, fluorescence-activated cell sorting; wpt, weeks post transplantation

### Aggravated pathological severity in RTKO mice and their reduction in low-dose PBMC hu-mice

In the histopathological analysis to assess the severity of GvHD, prominent lesions were perivascular inflammatory cell aggregation; apoptosis; and atrophy in the liver, lung, and kidney. In the skin, sporadic inflammatory cell infiltration, mild epidermal hyperplasia, and hyperkeratosis were observed (Fig 3A). The severity of these lesions ranged from moderate to severe in the lung and liver, whereas it was mild in the kidney and skin in the MD and HD groups. In the lower-dose PBMC groups, the severity of lesions and inflammatory cell aggregation was significantly reduced and minimal apoptosis and atrophy were observed in the lungs, liver, and kidneys. However, moderate epithelial thickening and hyperkeratosis were prominent pathological findings in the LD groups (Fig 3A). Overall, lesion scores were significantly higher in the RTKO-HD group and lower in the RID-LD group. Conversely, skin lesion scores were significantly higher in the LD groups, particularly in the RTKO-LD group (Fig 3B). No life-threatening lesions were observed, except in the liver and lung of the HD groups. Although the gut is recognized as one of the primary organs affected by GvHD [29], no abnormal lesions or leukocyte aggregations were detected in any of the groups (S3 Fig). Given that there were no significant differences in lesion scores between the LD-groups (Fig 3B), the GvHD symptoms observed in the RTKO-LD group were comparable to those in the RID-LD group.

**Fig 3.**
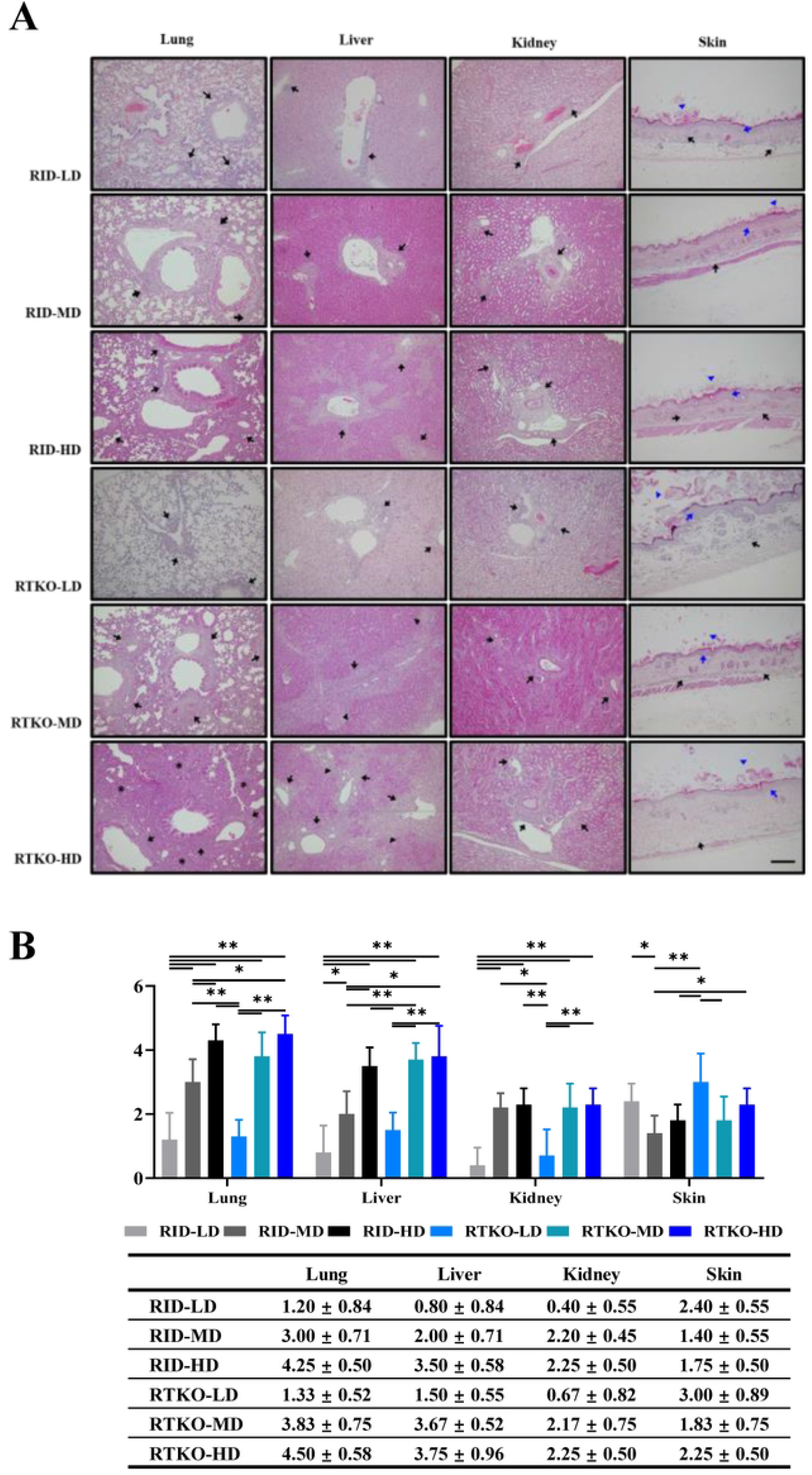
Histopathological findings and semi-quantitative lesion score analysis. (A) Representative hematoxylin and eosin (H&E)-stained sections of the lung, liver, kidney, and skin tissues. Prominent lesions are indicated as follows: black arrows, inflammatory cell aggregation; black arrowheads, necrosis with apoptosis; black asterisks, pulmonary consolidation; blue arrows, epidermal hyperplasia; blue arrowheads, hyperkeratosis. Scale bar = 100 µm. (B) Semi-quantitative lesion scores for lung, liver, kidney, and skin tissues. The RTKO-HD group exhibited the highest lesion scores across all evaluated organs. In the LD groups, the severity of lesions was significantly reduced, but skin lesions were aggravated. The number of mice in each experimental group was as follows: RID-LD (n = 5), RID-MD (n = 5), RID-HD (n = 4), RTKO-LD (n = 6), RTKO-MD (n = 6), RTKO-HD (n = 4). *P < 0.05. RID, Rag2; IL2rg double KO NOD mice; RTKO, CD47; Rag2; IL2rg triple KO NOD mice; LD, low dose; MD, middle dose; HD, high dose

### Comparative expression analysis of human leukocyte antigens in affected tissues and the spleen

Immunohistochemistry was performed to identify infiltrated human immune cells. hCD45+, hCD3+, and hCD19+ leukocytes were found to be aggregated in the perivascular areas and had infiltrated the interstitial tissues of the lung, liver, kidney, skin, and spleen. The majority of infiltrated human immune cells were hCD3+ T cells, accompanied by a smaller population of hCD19+ B cells (Fig 4A). For quantification, stained cells were manually counted within a 0.03 mm^2^ area under a light microscope. In the lungs, liver, kidneys, and spleen, the number of stained cells generally increased in a dose-dependent manner with PBMC administration, showing statistically significant differences. Conversely, the number of stained cells in the skin was significantly increased in the LD groups (Fig 4B). To elucidate the underlying cause of GvHD, human cytokines were measured using the Human CorPlex Cytokine Panel 1 10-Plex Array (Quanterix). Serum concentrations of hIL-4, hIL-12, hIL-22, and hTNFα were significantly increased in the RTKO groups compared to the RID groups. In contrast, hIL-5 levels were markedly decreased in the RTKO groups (Fig 5). No meaningful differences were observed for other cytokines, including hIL-1β, hIL-6, hIL-8, and hIFNγ; additional data for cytokines detectable across all groups are provided in S4 Fig.

**Fig 4.**
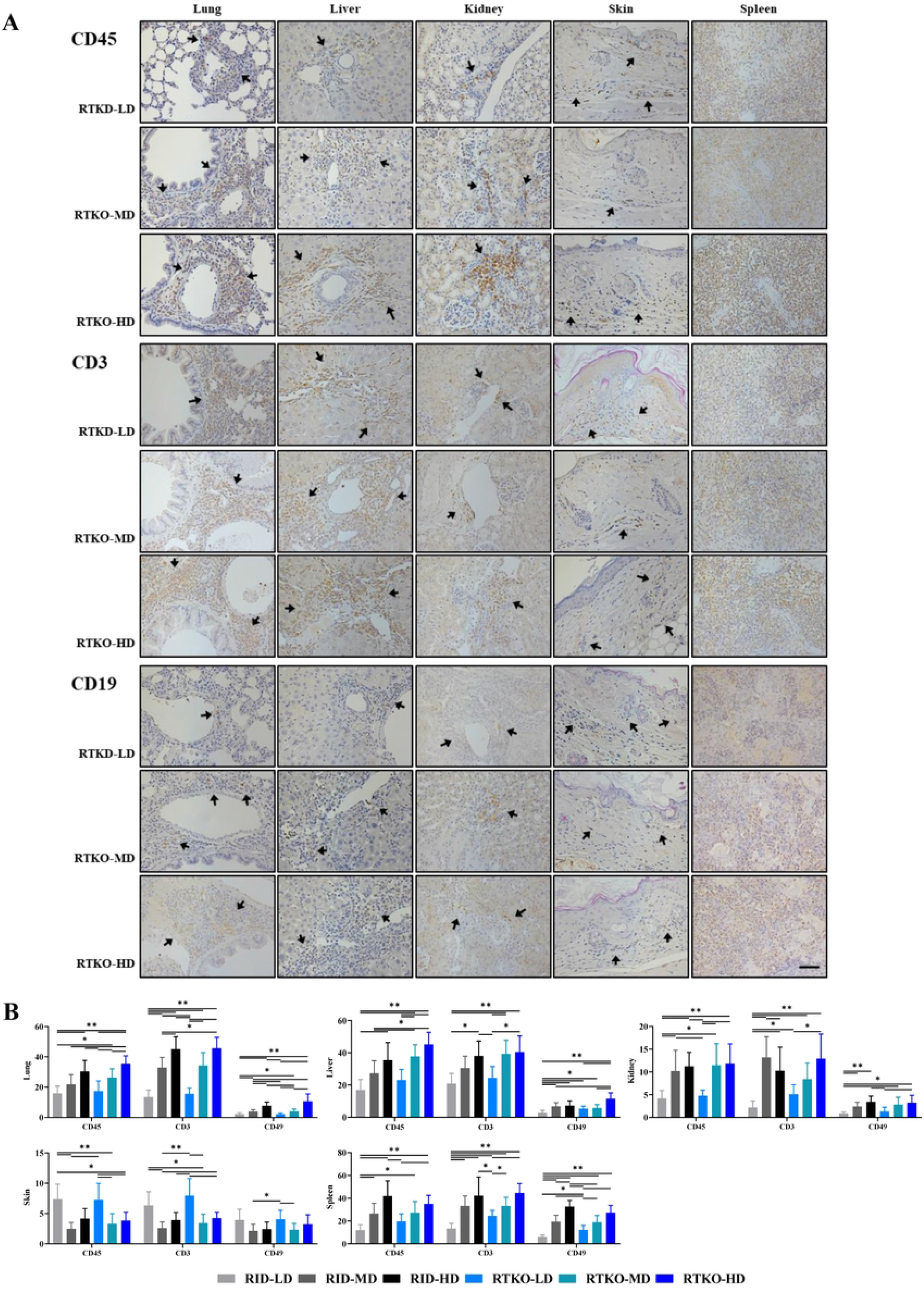
Immunohistochemical staining and quantification of humans leukocytes infiltration. (A) Representative IHC stains for hCD45, hCD3, and hCD19 in the lung, liver, kidney, skin, and spleen tissues of the RTKO groups. hCD45 and hCD3 were abundantly expressed in inflammatory cells, with a subset of leukocytes positive for hCD19. Stained regions or cells are indicated by arrows, except for the spleen, in which most cells were labelled. Stained cells increased in a PBMC dose-dependent manner and were higher in the RTKO groups compared to the RID groups. (B) Quantitative analysis of DAB-stained cells in the lung, liver, kidney, skin, and spleen. Positive cells were manually counted within a 0.03 mm^2^ area under a light microscope. Statistical analysis was performed based on counts obtained from three representative fields per slide. Scale bar = 50 µm. The number of mice in each experimental group was as follows: RID-LD (n = 5), RID-MD (n = 5), RTKO-LD (n = 6), RID-HD (n = 4), RTKO-MD (n = 6), RTKO-HD (n = 4). *P < 0.05 and **P < 0.01. h, human; RID, Rag2; IL2rg double KO NOD mice; RTKO, CD47; Rag2; IL2rg triple KO NOD mice; MD, middle dose; HD, high dose

**Figure 5.**
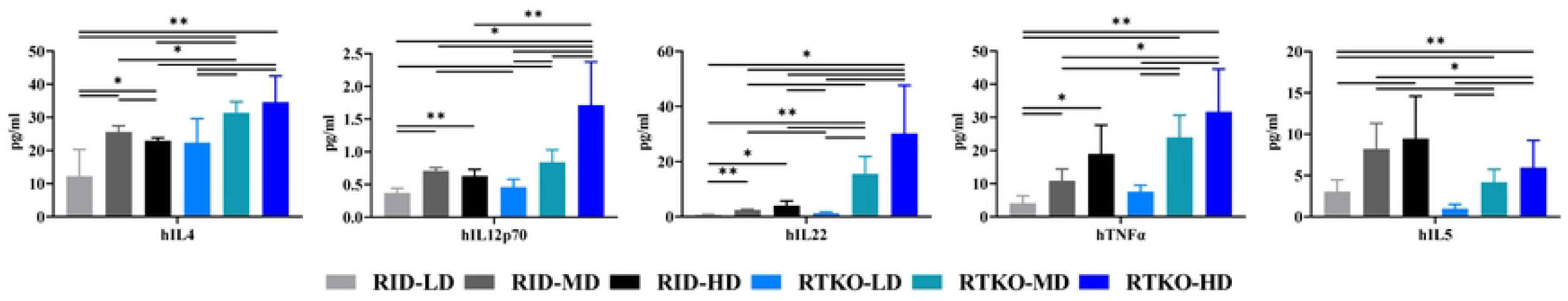
Serum levels of human cytokines analyzed via Multiplex ELISA. Serum cytokine concentrations were measured at the time of sacrifice. In the RTKO groups, the expression levels of hIL-4, hIL-12, hIL-22, and hTNFα increased, whereas that of hIL-5 decreased compared to the RID groups. The number of mice in each experimental group was as follows: RID-LD (n = 5), RID-MD (n = 5), RTKO-LD (n = 6), RID-HD (n = 4), RTKO-MD (n = 6), RTKO-HD (n = 4). *P < 0.05 and **P < 0.01. h, human; RID, Rag2; IL2rg double KO NOD mice; RTKO, CD47; Rag2; IL2rg triple KO NOD mice; MD, middle dose; HD, high dose; IL, interleukin; TNFα, tumor necrosis factor alpha; ELISA, enzyme-linked immunosorbent assay

## Discussion

Hu-mice models, which recapitulate the human immune system, are widely utilized as essential biological tools for various immunotherapy studies, including oncology, autoimmune diseases, and transplantation. PBMC- and HSC-engrafted hu-mice are the most commonly used models, each possessing distinct strengths and limitations [2, 3, 14]. To address these constraints, numerous studies have explored including the use of genetically engineered mouse strains [15, 16]. We previously developed RTKO mice [27] focusing on the strong affinity between SIRPα and human CD47 and functional immunodeficiencies of the NOD strain [11–13] and the enhanced tolerance of CD47 deficiency to support human leukocyte engraftment [16]. Using novel RTKO mice, we established an HSC hu-mice model that exhibited improved human leukocyte reconstitution and alleviated GvHD symptoms. Despite these advantages, a prolonged production period remained a major drawback; substantial leukocyte engraftment took eight weeks, and T-cell maturation necessitated more than twice as much time [27]. To overcome these limitations, the present study established the first RTKO-based PBMC humanized mouse model, aiming to achieve enhanced immune cell reconstitution and attenuated GvHD symptoms while maintaining the rapid production characteristic of PBMC models.

In the first trial, standard doses of PBMCs—5 × 10^6^ (MD) and 1 × 10^7^ (HD) cells—were administered to RID and RTKO mice [30, 31]. Although human immune cells were successfully engrafted in all groups, RTKO mice showed superior transplant efficiency, as indicated by the reconstitution of human leukocytes, including elevated hCD45 ratios (Fig 1). The RTKO-HD group initially had the highest engraftment of human leukocytes (37.7% of hCD45 at 2 wpt), but this declined rapidly starting at 4 wpt (Fig 1D), likely owing to GvHD symptoms. Considering weight loss, GvHD occurred from 3 wpt in the HD groups and from 4 wpt in the MD groups (Fig 1B), and severe symptoms such as jaundice, anemia were observe from 5wpt (Fig 1A, Table 1). The RTKO-MD group was considered to exhibit the best transplant efficiency, with the peak hCD45 ratio at 5 wpt (47.7%) and levels consistently maintained above 30% throughout the experimental period (Fig 1D). Animal fatalities began at 5 wpt in the HD groups and at 6 wpt in the MD groups (Fig 1A). Histopathological analysis revealed significant lesions in RTKO mice, although life-threatening lesions were only observed in the liver and lung of mice in the HD groups (Fig 3). Considering the reconstitution of human leukocytes and GvHD symptoms (Figs 1 and 3), the RTKO-MD group emerged as the most appropriate model in this trial. However, the occurrence of severe GvHD and the narrow experimental window of just six weeks were substantial drawbacks.

To address these limitations, we optimized the number of transplanted PBMCs, and developed a low-dose (3 × 10^6^) PBMC-engrafted hu-mice model. In the LD groups, the hCD45 ratio exceeded 5% at 4 wpt, indicating that the repopulation of human leukocytes was delayed compared to the MD and HD groups. Thereafter, the RTKO-LD group showed excellent transplant efficacy, with the hCD45 ratio maintained above 10% for 10 weeks from 5 wpt and above 20% for 7 weeks. While the health status of the RID-LD group was more stable, the ratio of human leukocytes remained below 10%, except between 6 and 8 wpt, and showed substantial individual variation (Fig 2C, S1 Table). Examining weight loss, GvHD generally began to occur at 13 wpt in the RID-LD group and at 11 wpt in RTKO-LD group (Fig 2B). However, some mice died or were euthanized before 10 wpt due to severe GvHD signs including jaundice, anemia, and hypothermia (Table 1). In the RTKO LD group, GvHD symptoms appeared to be alleviated, as there was no significant difference in GvHD severity between LD groups (Figs 2A, 2B, and 3B) despite the RTKO-LD group demonstrating significantly increased engraftment compared to the RID-LD group (Figs 2C–2F). Furthermore, while two out of three mice in the RID-LD group with hCD45^+^ levels exceeding 30% succumbed to GvHD, only two out of six mice in the RTKO-LD group died, suggesting that RTKO mice exhibit higher tolerance to human leukocyte transplantation (Fig 2A, S1 Table). Considering both transplant efficiency and health status, the RTKO-LD model provides a robust platform for stable, long-term experiments lasting approximately 10 weeks.

Compared to HSC hu-mice, PBMC hu-mice offer advantages such as a simpler generation procedure without requiring myelosuppression, easier access to human immune cells, and rapid reconstitution of the human immune system. However, a major drawback of conventional PBMC hu-mice is the narrow experimental window of just three to six weeks, necessitated by the onset of lethal GvHD [6, 32]. The significance of this study lies in the development of an advanced PBMC hu-mice model that addresses existing limitations and enables long-term experiments through the application of the novel RTKO strain and optimized PBMC dosing strategies. For HSC-humanized mice, while most experiments are concluded within 20 weeks post-transplantation, the optimal experimental window is typically around 8–10 weeks, considering the required timeframe for human immune cell differentiation [2, 32]. There are only a few reports of experiments extending beyond 12 weeks using HSC hu-mice [33]. Given that most tumor transplantation and immunotherapy studies are finalized within 10 weeks [6, 34], the extended experimental window provided by the RTKO-LD model is particularly noteworthy. Nevertheless, the poor reconstitution of immune cells other than T cells and B cells is a limitation of this model. The elevated hCD56 and hCD66b ratios observed in the RID group may suggest an abnormal immune response in one or two mice (Fig 1). Further research is required to achieve more consistent and robust reconstitution of diverse human immune cell populations.

Major necropsy findings in mice that died or were euthanized included inflammation and atrophy of the liver and lung. Histopathological analysis revealed severe inflammation in the liver and lung of the HD groups, and most of the infiltrated immune cells were hCD3+ T cells (Fig 4A). Serum levels of hIL-4, hIL-22, hIL-12, and hTNFα were increased in the RTKO groups, especially in the RTKO-HD group (Fig 5). hIL-12 is an inflammatory mediator in GvHD and plays a role in T cell proliferation [35], while hTNFα induces skin and liver damage and apoptosis as a core cytokine of acute and chronic GvHD [36, 37]. In addition, hIL-4 and hIL-22 aggravate skin, liver, and lung lesions [38, 39]. Most human serum cytokines are likely derived from T cells, which constitute the majority of human leukocytes. Consequently, the primary cause of GvHD in these models appears to be tissue damage caused by proliferating and infiltrating T cells and their associated cytokines. In the LD groups, cytokine expression was reduced compared to the MD and HD groups (Fig 5), which led to a significant reduction in the severity of the liver, lung, and kidney lesions (Figs 3A and 3B). However, the aggravated skin lesions observed in the LD groups (Fig 2) likely resulted from the prolonged influence of various cytokines over an experimental window that was eight weeks longer than that of the MD group. These skin lesions may have worsened due to the long-term effects of hIL-4 [40], hIL-22 [41, 42], hTNFα, and infiltrated leukocytes (Fig 4 and 5) involved in chronic cutaneous GvHD [14, 43]. hIL-5, a cytokine implicate in both acute and chronic GvHD pathogenesis, was decreased in the RTKO groups (Fig 5); this reduction may be linked to CD47 deficiency [41, 44]. We hypothesize that reduced hIL-5 levels may contribute to the alleviation of GvHD severity in RTKO mice, as evidenced by the comparable survival rates (Figs 1A and 2A) and the reduced severity of histopathological lesions (Fig 3) despite enhanced engraftment.

This study is the first to report the generation of PBMC humanized mice using CD47 deficient mice on a NOD strain, which offer significant immunological advantages for xenograft applications. Hu-mice models were successfully established across all experimental groups, using three different PBMCs transplantation doses in both RID and RTKO mice. Notably, RTKO mice exhibited significantly improved human leukocyte reconstitution and an extended experimental window, surpassing not only the RID controls but also conventional NSG/NRG models [42, 44]. Furthermore, the RTKO-LD model, with its optimized PBMC dosage, could serve as a robust platform for long-term, stable studies. By mitigating GvHD symptoms and extending the experimental window, it achieves a level of performance comparable to that of the HSC hu-mice model. Additionally, the radiation tolerance imparted by the Rag gene mutation enhances the suitability of this model for research in radiation oncology [7, 45]. We expect that the RTKO-LD PBMC hu-mice model developed in this study will serve as a remarkably versatile and valuable tool for various immunotherapy studies. However, further research is warranted to improve the repopulation of leukocytes other than T cells, reduce individual variation in allograft efficacy, and further alleviate GvHD symptoms.

## Supporting information

**S1 Fig. Immune monitoring of the humanized mouse applying standard doses of PBMC**. Engraftment of human cells was examined by FACS analysis. Representative dot plots of (A) hCD45, (B) hCD3 and hCD19, (C) hCD4 and hCD8, (D) hCD14 and hCD66b, and (E) hCD56 at 2 to 6 weeks after hPBMCs injection; FACS, fluorescence-activated cell sorting; RID, Rag2; IL-2rγ double KO NOD mice; RTKO, CD47; Rag2; IL-2rγ triple KO NOD mice; MD, middle dose; HD, high dose.

(TIF)

**S2 Fig. Immune monitoring of humanized mouse applying a lower dose of PBMC**. Engraftment of human cells was examined by FACS analysis 2 to 14 weeks after hPBMCs injection. Representative dot plots of hCD45, hCD3 and hCD19, hCD4 and hCD8, hCD14 and hCD66b in (A) RID and (B) RTKO mice; FACS, fluorescence-activated cell sorting; RID, Rag2; IL-2rγ double KO NOD mice; RTKO, CD47; Rag2; IL-2rγ triple KO NOD mice.

(TIF)

**S3 Fig. Hematoxylin and eosin staining of small and large intestines**. There was no abnormal lesion or aggregation of leukocyte in all groups. Scale bar = 100 µm. RID, Rag2; IL-2rγ double KO NOD mice; RTKO, CD47; Rag2; IL-2rγ triple KO NOD mice; LD, low dose; MD, middle dose; HD, high dose.

(TIF)

**S4 Fig. Measurement of human cytokines of the humanized mouse administered standard doses of PBMC**. Human cytokines were analyzed using Multiplex ELISA. Serum levels of (A) hIL-1β and (B) hIL-6. No significant differences were observed; RID, Rag2; IL-2rγ double KO NOD mice; RTKO, CD47; Rag2; IL-2rγ triple KO NOD mice; LD, low dose; MD, middle dose; HD, high dose.

(TIF)

**S1 Table. The individual hCD45 flow cytometry results of in the LD groups**.

(PDF)

## Acknowledgments

We thank the core facilities of the Department of Laboratory Animal Research and the Flowcytometry Core at the ConveRgence mEDIcine research cenTer (CREDIT), Asan Medical Center for the use of their shared equipment, services, and expertise. This work was also supported by the National Research Foundation of Korea (NRF) grant, funded by the Korean government (MSIT) (NRF-2020R1C1C1014653).

## Author contributions

**Conceptualization:** Sang-wook Lee, Seung-Ho Heo

**Data curation:** Kang-Hyun Kim, Sun-Min Seo

**Funding acquisition:** Seung-Ho Heo

**Investigation:** Kang-Hyun Kim, Hye-Young Song, Je-Won Ryu, Seon-Ju Jo, Seung-Hee Ryu, Sun-Min Seo

**Project administration:** Seung-Ho Heo

**Supervision:** Sang-wook Lee, Seung-Ho Heo

**Writing – original draft:** Kang-Hyun Kim, Hye-Young Song

**Writing – review & editing:** In-Jeoung Baek, Seung-Ho Heo

